# Primary mammary organoid model of lactation and involution

**DOI:** 10.1101/859645

**Authors:** Jakub Sumbal, Aurelie Chiche, Elsa Charifou, Zuzana Koledova, Han Li

**Author notes:** These authors contributed equally and should be considered co-first authors (Jakub Sumbal, Aurelie Chiche). Correspondence: Aurelie Chiche; Zuzana Koledova; Han Li.

## Abstract

Mammary gland development occurs mainly after birth and is composed of three successive stages: puberty, pregnancy and lactation, and involution. These developmental stages are associated with major tissue remodeling, including extensive changes in mammary epithelium as well as surrounding stroma. Three-dimensional (3D) mammary organoid culture has become an important tool in mammary gland biology and enabled invaluable discoveries on pubertal mammary branching morphogenesis and breast cancer. However, a suitable 3D organoid model recapitulating key aspects of lactation and involution has been missing. Here, we describe a robust and straightforward mouse mammary organoid system modeling lactation and involution-like process, which can be applied to study mechanisms of physiological mammary gland lactation and involution as well as pregnancy-associated breast cancer.

## Introduction

Lactation, the production of milk to feed progeny, is achieved by the mammary gland. This hallmark organ of mammals mainly develops postnatally and is highly dynamic (Macias and Hinck, 2012). With each pregnancy, mammary epithelium undergoes massive proliferation, tertiary branching of the mammary ductal system, and alveoli differentiation to prepare the epithelium for proper lactation (Brisken and Rajaram, 2006; Sternlicht, 2006). After parturition, mammary epithelium fully transforms into a milk-producing factory. Alveoli expand and take up space of regressing mammary stromal adipocytes, thereby multiplying epithelial volume many times (Macias and Hinck, 2012). After weaning, when milk production is no longer required, milk-producing epithelial cells are removed, and mammary gland is remodeled into a pre-pregnancy state. This process is called involution, which includes programed cell death of the epithelium, extracellular matrix (ECM) remodeling and re-differentiation of adipocytes (Hughes and Watson, 2012; Macias and Hinck, 2012; Zwick et al., 2018; Jena et al., 2019). By the end of involution, mammary gland is ready for a new cycle of pregnancy-associated growth, lactation and subsequent involution, which can be repeated throughout the reproductive lifespan. During these changes, mammary epithelium retains its bi-layered architecture with lumen facing luminal cells and basally situated myoepithelial cells, which is essential for proper function of the organ (Adriance et al., 2005; Haaksma et al., 2011; Macias and Hinck, 2012).

Endocrine signaling is a crucial regulator of mammary morphogenesis during pregnancy. Ovarian hormones estrogen and especially progesterone govern growth and morphogenesis of epithelium via induction of paracrine signaling between mammary stroma and epithelium, involving members of several growth factor families (Hennighausen and Robinson, 2005; Brisken and O’Malley, 2010). Pituitary hormone prolactin, on the other hand, acts directly on prolactin receptor on luminal cells and triggers alveoli maturation and lactogenic differentiation (Hennighausen and Robinson, 2005; Brisken and Rajaram, 2006). Involution is linked to cessation of hormonal stimuli and increase of inflammatory cytokines (Watson, 2006; Stein et al., 2007).

To study various aspects of mammary gland biology, three-dimensional (3D) cell culture models have been widely used for decades (Koledova, 2017a). They combine the advantages of easy manipulation of 2D cellular systems with providing complex cell-cell and cell-ECM interactions, thereby mimicking physiological conditions of in vivo experiments more faithfully (Shamir and Ewald, 2014; Huch and Koo, 2015; Koledova, 2017a; Artegiani and Clevers, 2018). Among the 3D culture models, primary mammary organoids have played a major role in understanding mechanisms of mammary branching morphogenesis (Ewald et al., 2008; Huebner et al., 2016; Neumann et al., 2018), including the role of ECM (Simian et al., 2001) and stromal cells (Koledova and Sumbal, 2019). Furthermore, spheroids produced from mammary cell lines were used to study tissue response to growth factors (Xian et al., 2005), organoids grown from sorted single primary mammary epithelial cells were used to study developmental potential of mammary epithelial cells (Linnemann et al., 2015;Jamieson et al., 2017), and differentiation of mammary-like organoids was achieved from induced pluripotent stem cells (Qu et al., 2017).

Despite these advances in 3D cell culture models of mammary gland, systems faithfully modeling pregnancy-associated morphogenesis and lactation have been spare. In some studies, β-casein or milk protein expression was used as a read-out of mammary epithelial functionality (Mroue et al., 2015; Jamieson et al., 2017). Several aspects of lactation and involution were captured in a co-culture of mammary epithelial and preadipocyte cell lines (Campbell et al., 2014) or in hormone-treated breast cancer cell spheroids (Ackland et al., 2003; Freestone et al., 2014). However, a system modeling lactation and involution in primary mammary organoids with proper architecture of bi-layered epithelium with myoepithelial cell layer was not characterized.

Here, we report on a mammary 3D culture system for studying induction and maintenance of lactation using easily accessible and physiologically relevant murine primary mammary organoids cultured in Matrigel. Upon prolactin stimulation, the organoids produce milk for at least 14 days and maintain a histologically normal architecture with a functional contractile myoepithelial layer. Moreover, upon prolactin signal withdrawal, our system recapitulates several aspects of involution. Altogether, we describe a robust, consistent and easy-to-do system for modeling crucial aspects of pregnancy-associated mammary gland morphogenesis and lactation.

## Materials and Methods

### Isolation of primary mammary epithelial organoids

Primary mammary organoids were isolated from 7-10 weeks old female mice (ICR or C57/BL6) as previously described (Koledova, 2017b). The animals were obtained from the Central Animal Facility of the Institut Pasteur and the Laboratory Animal Breeding and Experimental Facility of the Faculty of Medicine, Masaryk University. Experiments involving animals were approved in accordance with French legislation in compliance with European Communities Council Directives (A 75-15-01-3), the regulations of Institut Pasteur Animal Care Committees (CETEA), the Ministry of Agriculture of the Czech Republic, and the Expert Committee for Laboratory Animal Welfare at the Faculty of Medicine, Masaryk University. The study was performed by certified individuals (AC, JS, EC, ZK) and carried out in accordance with the principles of the Basel Declaration.

Briefly, the mice were euthanized by cervical dislocation, the mammary glands were removed, mechanically disintegrated and partially digested in a solution of collagenase and trypsin [2 mg/ml collagenase (Roche or Sigma), 2 mg/ml trypsin (Dutcher Dominique or Sigma), 5 μg/ml insulin (Sigma), 50 μg/ml gentamicin (Sigma), 5% fetal bovine serum (FBS; Hyclone/GE Healthcare) in DMEM/F12 (Thermo Fisher Scientific)] for 30 min at 37°C with shaking at 100 rpm. Resulting tissue suspension was treated with 20 U/ml DNase I (Sigma) and 0.5 mg/ml dispase II (Roche) and exposed to five rounds of differential centrifugation at 450 × g for 10 s, which resulted in separation of epithelial (organoid) and stromal fractions. The organoids were resuspended in basal organoid medium [BOM; 1× ITS, 100 U/ml of penicillin, and 100 μg/ml of streptomycin in DMEM/F12 (all from Thermo Fisher Scientific)] and kept on ice until used for 3D culture.

### 3D culture of mammary organoids

Freshly isolated primary mammary organoids were mixed with growth factor reduced Matrigel (Corning) and plated in domes. After setting the Matrigel for 45-60 min at 37°C, the cultures were overlaid with cell culture medium according to the experiment and incubated at 37°C, 5% CO2 in humidified atmosphere. The media used were: FGF2 medium [2.5 nM FGF2 (Peprotech or Thermo Fisher Scientific) in BOM]; Lactation medium [LM; 1 μg/ml prolactin (mouse recombinant for qPCR, western blot, immunohistochemistry and contraction control; Sigma or Peprotech) or sheep pituitary prolactin for time-lapse and confocal imaging, and contraction experiments; Sigma) and 1 μg/ml hydrocortisone (Sigma) in BOM]. FGF2 medium was changed every 3 days, LM was changed every 2 days.

For time-lapse imaging experiments, organoids were incubated in humidified atmosphere of 5% CO2 at 37°C on Olympus IX81 microscope equipped with Hamamatsu camera and CellR system for time-lapse imaging. For morphological analysis of organoid development, the organoids were photographed from day 8 to day 17 of culture, one image per organoid was taken every hour. The images were exported and analyzed using Image J (NIH). For analysis of organoid contraction, the organoids were photographed from day 6 to day 20 of culture. On each imaging day, the pictures were taken every second for 120 seconds. The images were exported to video at 10 frames per second using xCellence software (Olympus).

### Whole mount staining of mammary organoids

Organoid cultures were fixed with 10% neutral buffered formalin for 30 min and washed twice with PBS. Then, the cultures were rinsed with 70% ethanol and stained with Oil Red O solution (Koopman et al., 2001; Kim et al., 2015) [0.3% (w/v) Oil Red O (Sigma) in 70% (v/v) ethanol] for 30 min in the dark. Subsequently, the cultures were washed with 70% ethanol for 10 min and twice with PBS, 5 min each wash. Then the cultures were stained with 0.5 μg/ml DAPI and 2 U/sample phalloidin-AlexaFluor488 (Thermo Fisher Scientific) in PBS for 1 h at room temperature (RT) in the dark. After two washes with PBS, the cultures were attached to a coverslip-bottom dish (ibidi) with 1% low gelling temperature agarose (Sigma) and overlaid with PBS. The organoids were imaged using an LSM800 confocal microscope (Zeiss) with and analyzed using ZEN blue software (Zeiss).

### Histology and immunostaining analysis

For histological analysis, organoids were washed 3 times with PBS and fixed for 30 min in 4% paraformaldehyde (Electron Microscopy Sciences). After washing with PBS, organoid 3D cultures were embedded in 3% low gelling temperature agarose. After solidification, the samples were dehydrated and embedded in paraffin. Sections (5 μm thick) were cut and dewaxed for hematoxylin and eosin staining or immunostaining. For localization of prolactin receptor expressing cells, 10 µm cryosections of mammary glands from *Prlr-IRES-Cre;ROSA26-CAGS-GFP* mice (Aoki et al., 2019) were labeled with antibodies and counterstained with 0.5 μg/ml DAPI, mounted with Vectashield (Vector Labs) and images were taken on LSM800 microscope (Zeiss).

The following primary antibodies were used: Goat anti-GFP (Origene; R1091P; 1:200), rabbit polyclonal anti-keratin 5 (BioLegend, 905501; 1:200), mouse monoclonal anti-keratin 8 (BioLegend, 904801; 1:200), and mouse monoclonal anti-β-casein (Santa Cruz, sc-166530; 1:250). Corresponding secondary antibodies were used: Donkey anti-rabbit Dylight 488 and donkey anti-mouse Dylight 594 (Immuno Reagents; DkxRb-003-D594NHSX and DkxMu-003-D488NHSX; 1:200), together with 1 μg/ml of Hoechst-33342 (Thermo Fisher Scientific) for immunofluorescence labeling, or anti-mouse HRP associated secondary antibody (Dako).

### RNA isolation and real-time quantitative PCR

Total RNA was extracted from organoid samples using RNeasy Micro Kit (Qiagen) following manufacturer’s instructions. Reverse transcription was performed using High capacity cDNA reverse transcription kit (Thermo Fisher Scientific). Quantitative real-time PCR was performed using 5 ng cDNA, 5 pmol of the forward and reverse gene-specific primers each in Light Cycler SYBR Green I Master mix (Roche) on LightCycler 480 II (Roche). All reactions were performed at least in duplicates and in a total of at least two independent assays. Relative gene expression was calculated using the ΔΔCt method and the values were normalized to housekeeping gene *Gapdh*. The primers of following sequences (5’-3’) were used: *Csn2*-forward (F): CCTCTGAGACTGATAGTATTT, *Csn2*-reverse (R): TGGATGCTGGAGTGAACTTTA; *Wap*-F: TTGAGGGCACAGAGTGTATC, *Wap*-R: TTTGCGGGTCCTACCACAG; *Mmp3*-F: CCTGATGTTGGTGGCTTCA, *Mmp3*-R: TCCTGTAGGTGATGTGGGATTTC; *Mmp13*-F: ACTTCTACCCATTTGATGGACCTT, *Mmp13*-R: AAGCTCATGGGCAGCAACA; *Gapdh*-F: TTCACCACCATGGAGAAGGC, *Gapdh*-R: CCCTTTTGGCTCCACCCT. All primers were purchased from Sigma.

### Western blot

3D cultures were dissociated by repetitive pipetting in ice cold PBS with phosphatase inhibitors (Merck) and centrifuged (450 × g, 3 min at 4°C). Supernatant containing Matrigel proteins was discarded and pellets were lyzed in RIPA buffer (Merck), supplied with protease and phosphatase inhibitors (Merck). After vortexing and sonication, protein lysates were cleared by centrifugation and protein concentration was measured using Coomassie reagent (Merck). Denatured, reduced samples were resolved on 12.5% SDS-PAGE gels (Bio-Rad) and blotted onto nitrocellulose membranes by Trans-blot Turbo transfer system (Bio-Rad). After blotting, the membranes were blocked with 2% BSA in PBS with 0.1% Tween-20 (Merck; blocking buffer) and incubated with primary antibodies diluted in blocking buffer overnight at 4°C. After washing in PBS with 0.05% Tween-20, the membranes were incubated with horseradish peroxidase-conjugated secondary antibodies for 1 h at RT. Signal was developed using an ECL substrate (Thermo Fisher Scientific) and imaged with ChemiDoc MP imaging system (Bio-Rad) and band density was analyzed in ImageJ.

The following antibodies were used for immunoblotting: Mouse monoclonal anti-β-casein (Santa Cruz, sc-166530, 1:1000), mouse monoclonal anti-α-tubulin (Santa Cruz, sc-5286, 1:1000), and anti-mouse secondary antibody (Merck, NA931, 1:1000).

### Statistical analysis

Statistical analysis was performed using the GraphPad Prism software, statistical test used is specified in figure legend. * p < 0.05, ** p < 0.01, *** p < 0.001, **** p < 0.0001. The number of independent biological replicates is indicated as n.

## Results

### FGF2 pre-treatment enhances lactogenic differentiation of mammary epithelium

During mammary gland morphogenesis, lactation is preceded by excessive branching of epithelial ducts. Therefore, we tested if epithelial expansion by branching morphogenesis is required for lactogenic differentiation *in vitro*. To this end, the primary mammary epithelial organoids were either treated with lactation medium (LM, containing prolactin and hydrocortisone) only for 4 days; or treated with FGF2, a potent mammary epithelium branching-inducing factor (Ewald et al. 2008), for 6 days and followed by 4 days of LM (Fig. 1A, S1A). To detect lactogenic differentiation, we measured the expression of *Csn2* and *Wap* by RT-qPCR.

**Figure 1.**
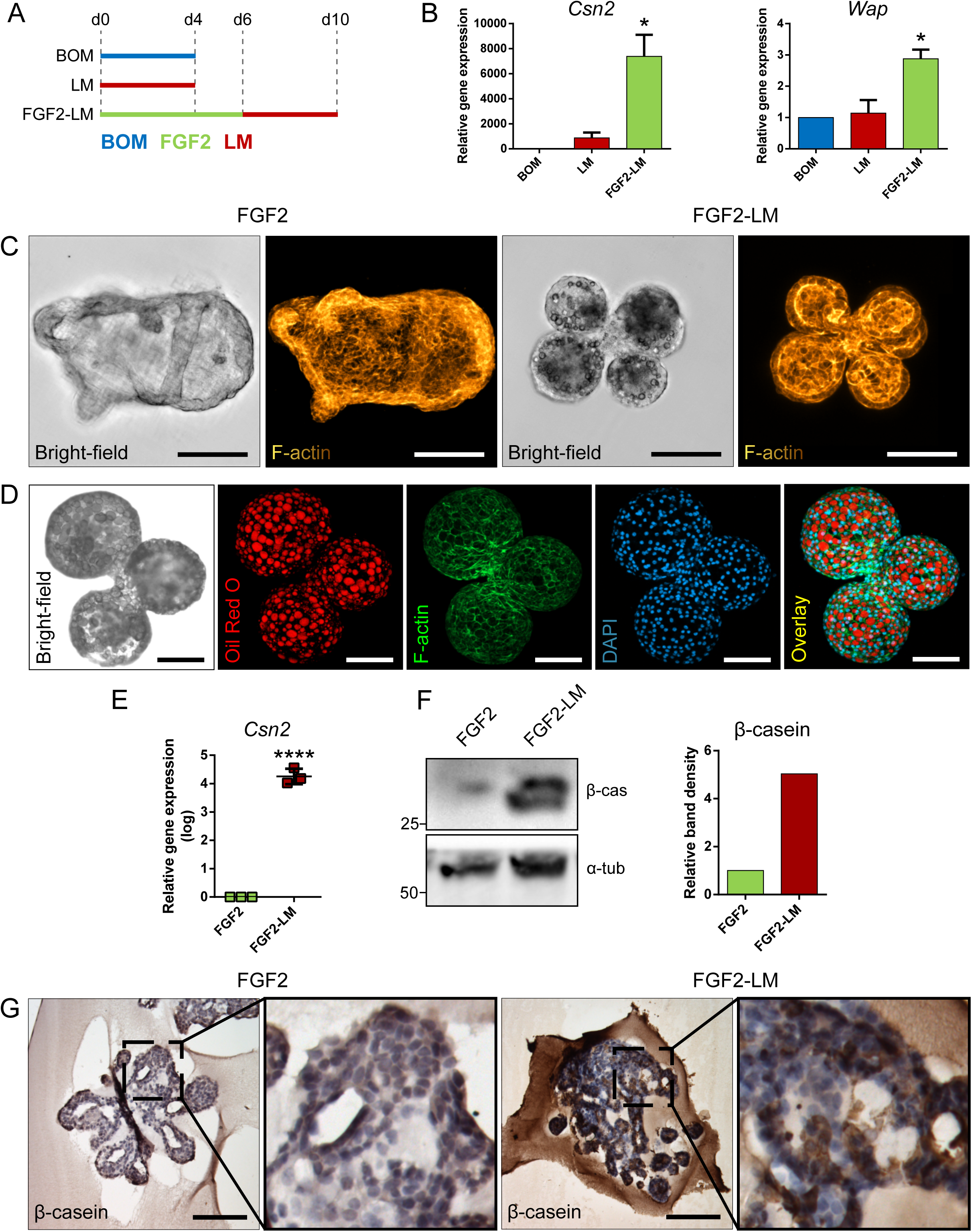
Lactation induction in primary mammary organoids. (A, B) FGF2 pre-treatment increases lactation capacity of primary mammary organoids. (A) Scheme depicting the experimental design. BOM, basal organoid medium; LM, lactation medium; FGF2, FGF2 medium. (B) Expression of milk genes *Csn2* and *Wap* in organoids treated with BOM, LM, or FGF2 followed by LM. The plot shows mean + SD; n = 2. One-way ANOVA, * p < 0.05. (C) Bright-field images and maximum intensity projection images from confocal imaging of whole-mount organoids after treatment with FGF2 only or with FGF2 followed by LM (Yellow-to-brown staining shows F-actin). Scale bars represent 100 μm. (D) Bright-field image and maximum intensity projection images from confocal imaging of whole-mount organoid treated with FGF2 followed by LM. Red, Oil Red O (lipids); green, F-actin; blue, DAPI (nuclei). Scale bars represent 100 μm. (E) Immunohistochemical staining of β-casein in organoids treated with FGF2, or FGF2 and then LM. Marked area is shown in higher magnification. Scale bars represent 100 μm. (F, G) Quantification of β-casein expression in organoids treated with FGF2, or FGF2 followed by LM. (F) RT-qPCR analysis of β-casein gene Csn2. The plot shows mean + SD; n = 3. Unpaired Student’s t-test, two tailed, **** p < 0.0001. (G) Western blot analysis of β-casein expression on protein level. The plot shows quantification of band density.

Our results revealed that treatment of freshly isolated organoids with LM only induced the expression of *Csn2* (Fig. 1B). However, when organoids were pre-treated with FGF2, the expression of both *Csn2* and *Wap* were significantly increased (Fig. 1B). These data suggest the branching morphogenesis induced mammary epithelial expansion could enhance the lactogenic ability of mammary epithelium.

### Lactation medium induces production of milk proteins and secretion of lipid droplets

Next, we compared the morphology of organoids treated either with FGF2 only or FGF2 and LM (FGF2-LM) to further characterize the phenotype of lactation organoids. On bright-field micrographs, we noticed that FGF2-LM organoids appeared to have a darker lumen, possibly due to the milk accumulation (Fig. 1C). Interestingly, we also observed bubble-like structures at the apical site of epithelium in the same organoids, which potentially represented lipid droplets (Fig. 1C). To further characterize these droplets, we stained the organoids with F-actin (with phalloidin), a cytoskeletal protein, or with Oil Red O. Confocal microscopy revealed that the droplets were negative for F-actin and strongly positive for Oil Red O, confirming the droplets are lipid (Fig. 1C, D). Next, we assessed the expression of milk proteins in the organoids. First, we detected a significant increase in *Csn2* by four orders in FGF2-LM treated organoids compared to FGF2 alone by RT-qPCR (Fig. 1E). Consistently, in FGF2-LM treated organoids, we detected upregulation of β-casein on the protein level by Western blot (Fig. 1F) and a strong cytoplasmic signal by immunohistochemistry (Fig. 1G).

Taken together, these data demonstrate that mammary primary organoids are capable of milk production after prolactin treatment, which could be greatly enhanced by branching morphogenesis.

### Morphology maintenance in long-term lactating organoids

After successful induction of lactation in the primary mammary organoids with the FGF2-LM protocol, we went on to investigate the lactation-associated phenotype in long-term organoid culture. After 6 days FGF2 treatment, the organoids were cultured either continuously with LM (FGF2-LM) or switched to BOM after 4 days LM treatment (FGF2-LM-BOM) (Fig. 2A). The morphogenesis of the organoids was recorded using time-lapse microscopy for 20 days. Interestingly, FGF2-LM-BOM cultured organoids regressed both in size and the complexity of the shape, while the organoids in FGF2-LM maintained the size and only partially lost the branched phenotype (Fig. 2B, C). In addition, unlike the organoids in FGF2-LM-BOM, the organoids in FGF2-LM retained the darker appearance, possibly due to the milk accumulation (Fig. 2B, D).

**Figure 2.**
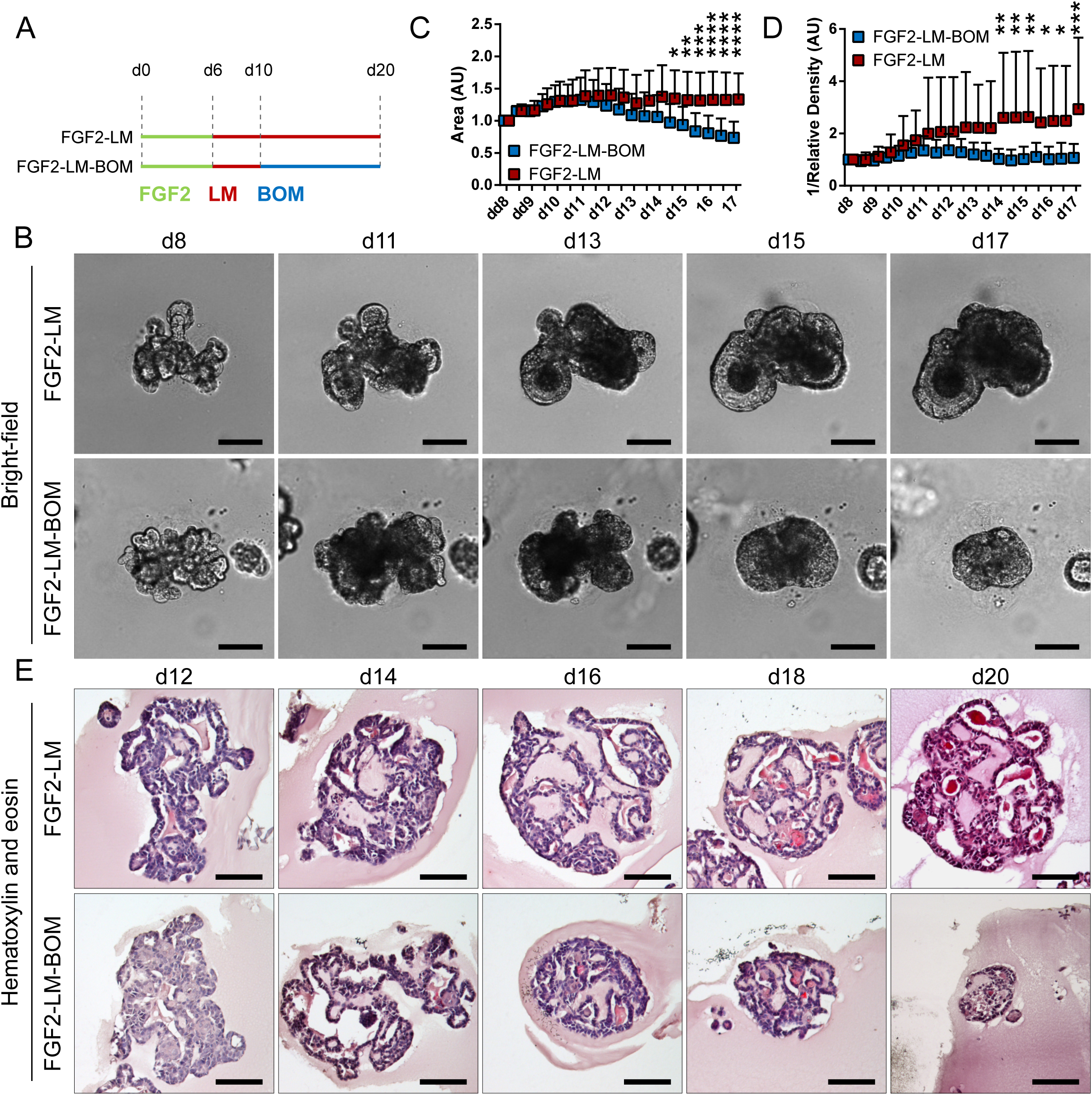
Morphology of organoids undergoing long-term lactation. (A) Scheme depicting experimental design. FGF2, FGF2 medium; LM, lactation medium; BOM, basal organoid medium. (B) Bright-field images from time-lapse imaging of organoid morphogenesis under continuous LM treatment or under LM withdrawal and replacement with BOM (LM-BOM). Scale bars represent 100 μm. (C, D) Morphometric analysis of organoid area (C) and density (D) from the time-lapse experiment. The plots show mean + SD; n = 2, N = 20 organoids per condition. Two-way ANOVA, * p < 0.05, ** p < 0.01, *** p < 0.001, **** p < 0.0001. (E) Hematoxylin and eosin staining of organoids at different time-points of long-term lactation. Scale bars represent 100 μm.

Morphologically, FGF2-LM treated organoids exhibited complex architecture with multiple lumens filled with dense eosinophilic material, which was maintained throughout the experiment (Fig. 2E, upper panel). However, upon LM withdrawn, the complex architecture was lost rapidly and organoids involuted into small spheroid with much simpler structures (Fig. 2E, lower panel).

### Milk production in long-term lactating organoids

Of note, we detected strong β-casein signal in the intraluminal of long-term lactating organoids by immunohistochemistry. Closer observation revealed that cytoplasmic β-casein signal was sustained in long-term LM culture (Fig. 3A, upper panel), but lost after LM withdrawal (Fig. 3A, lower panel). In addition, RT-qPCR revealed that FGF2-LM treated organoids maintained high level of *Csn2* expression, which was dramatically reduced by 4 to 5 orders of magnitude in FGF2-LM-BOM treated organoids (Fig. 3B). The same change was confirmed in the protein level by Western blot (Fig. 3C). Therefore, the production of β-casein depends on the prolactin signaling. Altogether, these data suggest that these organoids have proper epithelial architecture and the capacity to maintain milk production over prolonged culture time in response to the prolactin signaling.

**Figure 3.**
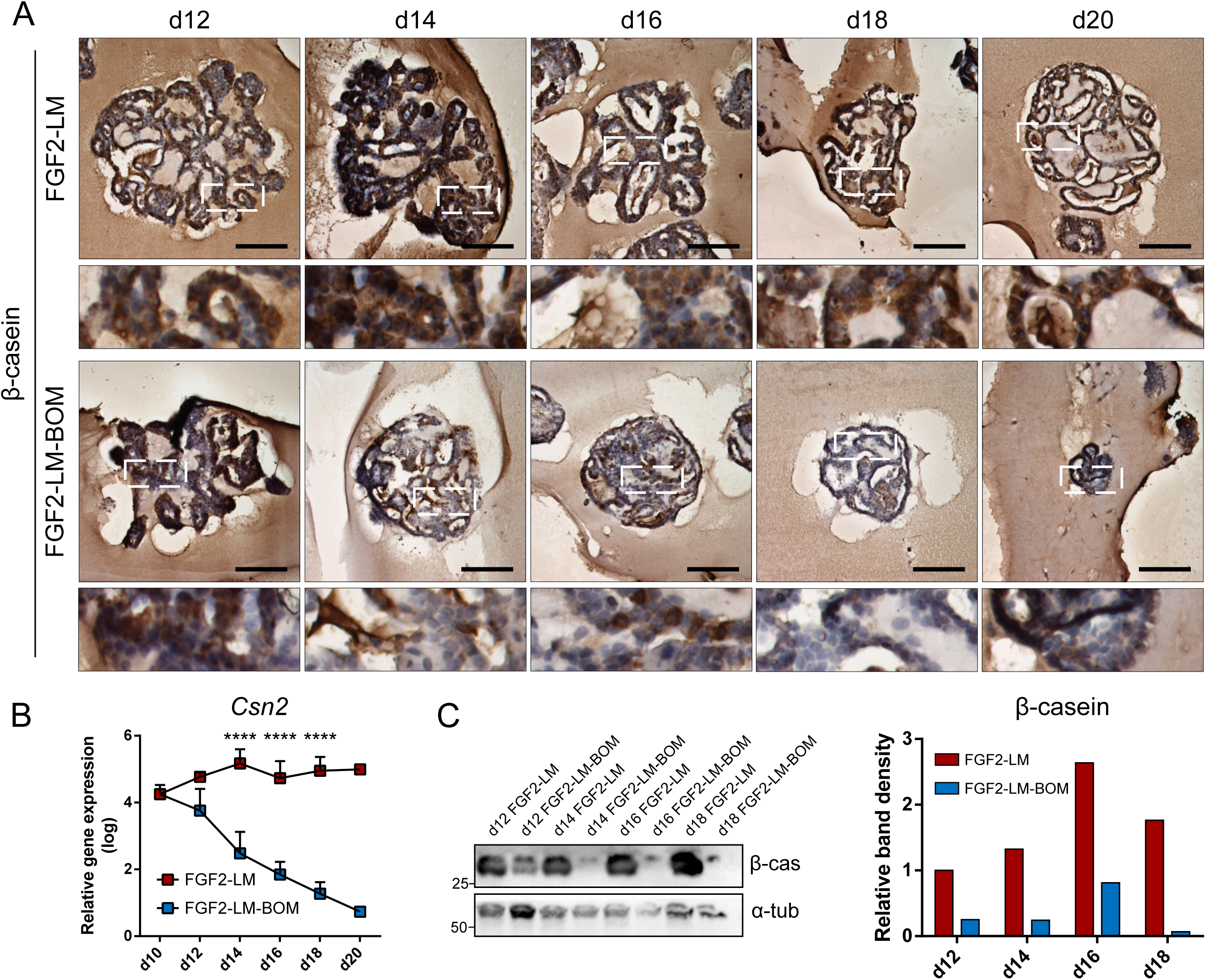
Milk production during long-term lactation. (A) Immunohistochemical staining of β-casein in organoids during long-term LM treatment or after LM withdrawal (LM-BOM), according to experimental scheme in Fig. 2A. Marked area is shown in higher magnification. Scale bars represent 100 μm. (B) *Csn2* expression during long-term lactation with continuous lactation medium (LM) or with hormonal withdrawal (LM-BOM). The plot shows mean + SD; n = 3 for d12 to d18, n = 1 for d20. Two-way ANOVA, **** p < 0.0001. (C) Western blot analysis and band density quantification of β-casein expression in organoids during long-term lactation.

### Lactating organoids retain functional myoepithelial layer with contractility

Next, we co-stained the lactating organoids with keratin 5 and keratin 8, markers of myoepithelial and luminal cells respectively, to confirm that the organoids contain proper bilayer epithelial architecture. We found that FGF2-LM treated organoids contained a continuous layer of myoepithelial cells, similar to FGF2 treated organoids (Fig. 4A). Moreover, the myoepithelial cell layer was retained during the long-term culture in LM treatment as well as after LM withdrawal (Fig. 4B), suggesting the luminal-myoepithelial cell homeostasis was stable during long-term culture.

**Figure 4.**
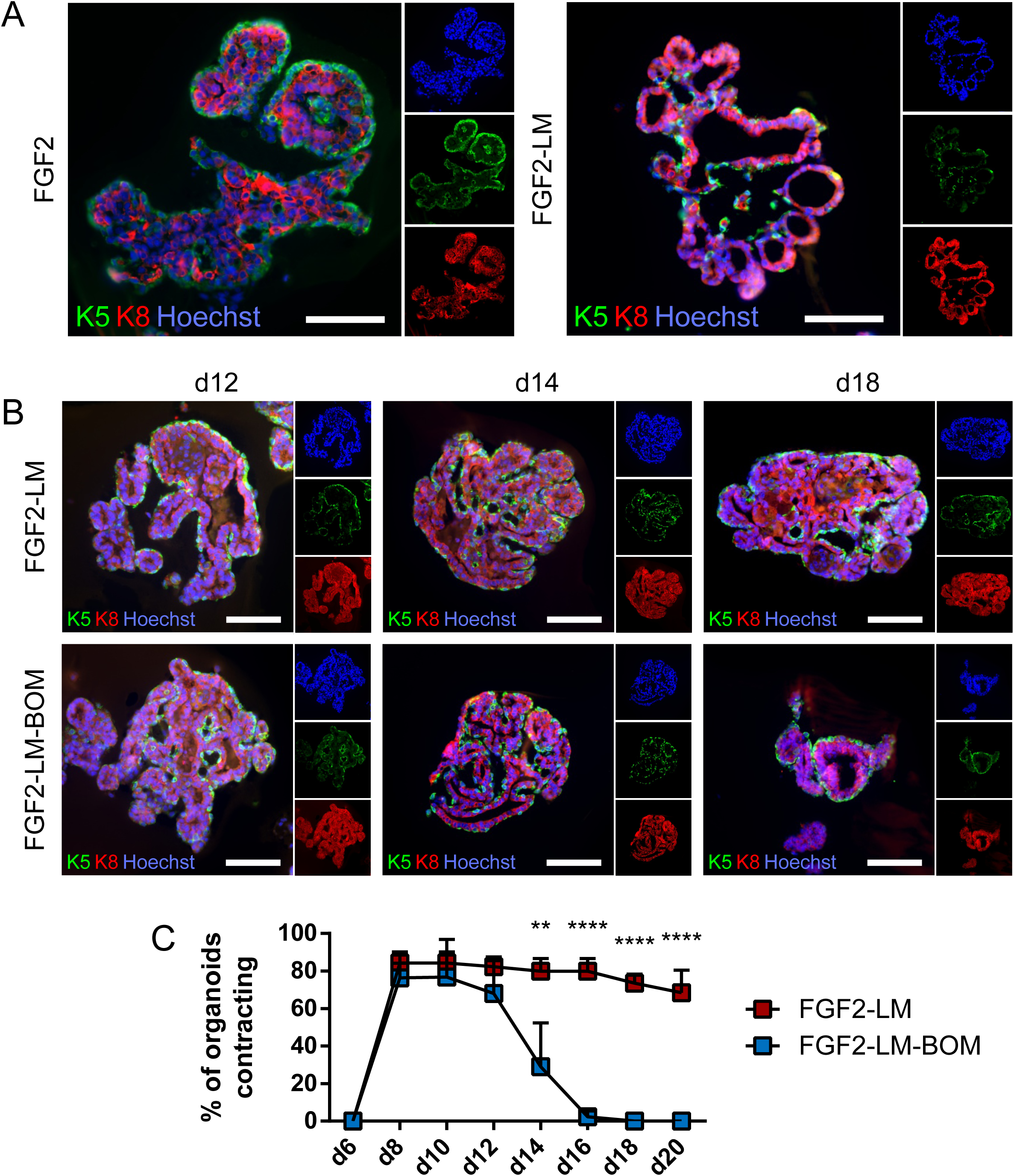
Lactating organoids retain functional myoepithelial layer. (A) Immunofluorescent staining shows distribution of myoepithelial (keratin 5 positive, green) and luminal cells (keratin 8 positive, red) in organoids treated with FGF2 or FGF2 followed by LM. Hoechst, blue (nuclei). Scale bars represent 100 μm. (B) Immunofluorescent staining shows distribution of myoepithelial (keratin 5 positive, green) and luminal cells (keratin 8 positive, red) in organoids during long-term lactation. Hoechst, blue (nuclei). Scale bars represent 100 μm. (C) Quantification of contracting organoids from movies recorded at indicated time-points. The plot shows mean + SD; n = 2, N = 50 organoids per experiment. Two-way ANOVA, ** p < 0.01, **** p < 0.0001.

Importantly, FGF2 treatment induced stratification of the luminal layer, which is in agreement with published work (Fig. 4A) (Ewald et al., 2008). Upon LM treatment, the organoids showed resolution of the stratified epithelium to predominantly bilayer structure, with luminal cells (keratin 8 positive) lining the luminal space (Fig. 4A, B), which is important for producing milk. Remarkably, we observed the LM treated organoids could contract periodically (Movie 2). In comparison, organoids never treated with LM showed relatively static structures (Movie 1). Of note, the contracting phenotype maintained during the long-term LM treatment, and quickly ceased after LM withdrawal (Fig. 4C). This result is somewhat puzzling since prolactin receptor is present only in the luminal cells (Fig. S2A). Of note, the prolactin used in our contraction experiments were isolated from sheep pituitary, which contains oxytocin (Vorherr et al., 1978). To test whether the contraction of myoepithelial cells is a direct effect of prolactin signaling, we compared contraction induction upon LM containing either sheep pituitary prolactin or mouse recombinant prolactin. Interestingly, that only sheep pituitary prolactin induced organoid contraction (Fig. S2B), suggesting that oxytocin contamination caused myoepithelial cell contraction.

These results demonstrate that myoepithelial layer is present in the lactating organoids. More importantly, these myoepithelial cells could contract in response to LM treatment, suggesting they are functionally similar to the *in vivo* counterpart.

### LM withdrawal triggers involution-like phenotype in lactating organoids

Involution is characterized by the regression of the lactating epithelium and remodeling of the mammary gland, which is induced upon the weaning of the pups (Jena et al., 2019). Interestingly, withdrawal of LM from lactating organoids also induced the size regression and loss of the branched morphology with luminal architecture (Fig. 2B-E). The cessation of milk production and ECM remodeling are the other two hallmarks of involution. Similarly, we detected the reduced β-casein signal (Fig. 3A, C) and *Csn2* expression (Fig. 3B) in the organoids upon LM withdrawal. Interestingly, we also found the expression of *Mmp2* and *Mmp13*, two important Mmps for the ECM remodelling process during involution, was upregulated in organoids after LM withdrawal (Fig. 5A, B).

**Figure 5.**
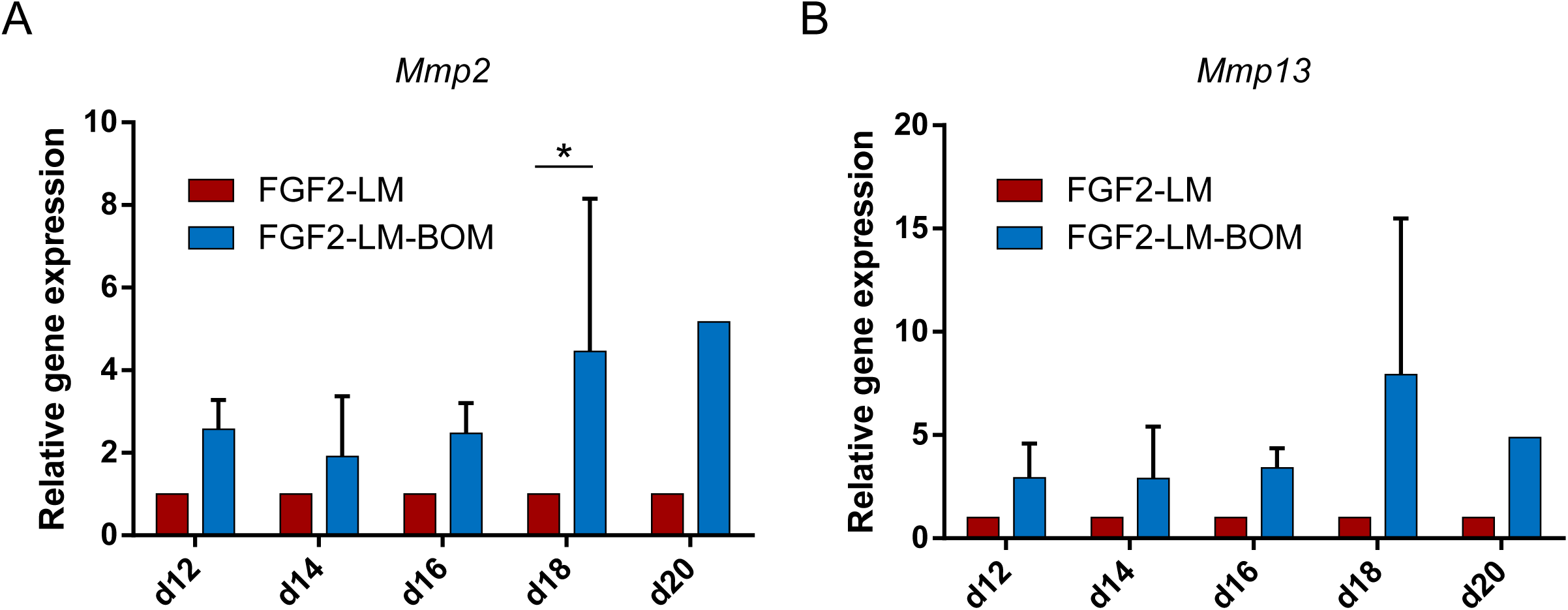
Expression of matrix metalloproteinases increases after lactation medium withdrawal. (A, B) RT-qPCR analysis of *Mmp2* (A) and *Mmp13* (B) expression in organoids during long-term lactation with continuous lactation medium (LM) treatment or with hormonal/LM withdrawal (LM-BOM). The plots show mean + SD; n = 3 for d12-d18, n = 1 for d20. Two-way ANOVA, * p < 0.05.

Together, these results demonstrate that upon withdrawal of hormonal stimulation, lactating organoids stop milk production and enter an involution-like process, thereby mimicking the *in vivo* situation upon weaning.

## Discussion

In this work, we described the use of primary mammary epithelial organoids to model pregnancy-associated morphogenesis and lactation. In our 3D culture system, primary mammary organoids exposed to LM with prolactin recapitulated several aspects of lactation process. Upon LM withdrawn, organoids regressed in a manner similar to the involution process *in vivo*.

Our data showed that FGF2 primes mammary epithelium for lactation. This is consistent with in vivo studies that noted morphological abnormalities in pregnancy-associated tertiary branching of mammary epithelium with attenuated FGF receptor signaling (Lu et al., 2008; Parsa et al., 2008). However, it remains to be elucidated what of the FGF2-mediated processes, including epithelial expansion, branching, and maturation, are essential contributors to milk production efficiency.

While several previous studies reported lactation induction in mammary epithelial organoids in response to prolactin in vitro, they did so only at a single time-point (Mroue et al., 2015; Jamieson et al., 2017). Long-term lactation in organoid cultures has not been reported before. In this study we documented milk production maintenance and stable morphology of lactating organoids over 14 days long lactation culture period. Physiological lactation in mouse lasts for circa three weeks (König and Markl, 1987) and milk composition and production rate vary during the lactation period to accommodate the needs of the offspring (Knight et al., 1986). We propose that our model would be suitable to study factors that influence dynamic changes in milk composition and quantity in long term.

Moreover, while previous studies used sample-destructive methods to detect lactation, such as organoid fixation and immunodetection of milk proteins (Mroue et al., 2015; Jamieson et al., 2017), we propose approaches for observing changes in milk production in the same organoid over time. They include morphological changes accompanying lactation in organoids that are easily observable by light microscopy and traceable by time-lapse imaging, namely appearance of lipid droplets in luminal space, increase in organoid darkness (integrated density) and the intriguing contraction of myoepithelial cells.

Myoepithelial cells form a layer of mammary epithelium that is situated basally to the luminal cells (Macias and Hinck, 2012). Together with their more recently elucidated role in keeping epithelial homeostasis and integrity (Adriance et al., 2005; Goodwin and Nelson, 2018; Sirka et al., 2018), their key function is to enable milk ejection by their contraction when pups are suckling (Haaksma et al., 2011). In response to tactile stimuli, oxytocin is released from pituitary and it binds to oxytocin receptor on myoepithelial cell to induce contraction (Nishimori et al., 1996; Froemke and Carcea, 2017). Therefore, oxytocin was employed to induce myoepithelial contraction in single cells (Raymond et al., 2011), as well as in an organoid system (Mroue et al., 2015). However, organoid contraction was shown only as decrease in organoid area over 20 minutes (Mroue et al., 2015). In contrary, we observed that contraction of a lactating organoid is a very fast process and the dynamic changes in organoid shape and size are visible to human eye. From videos of contracting organoids, recorded at the rate of one frame per second, we calculated that the frequency is about one contraction per ten seconds, which is very similar to the recently reported alveoli warping frequency of lactating mammary tissue upon oxytocin stimulation (Stewart et al., 2019). Although organoids in our study contracted without additional oxytocin treatment, the sheep pituitary prolactin used in our protocol contains oxytocin, which is likely the inducer of contraction, since mouse recombinant prolactin did not induce organoid contraction. Nevertheless, contracting organoids indicate that a functional myoepithelial layer is retained during the long-term lactation protocol.

Upon LM withdrawal, lactating organoids underwent involution-like changes: They regressed is size and complexity, and increased expression of MMPs, the proteases typically found in mammary gland during involution (Lund et al., 1996; Green and Lund, 2005). Involution-like morphological changes upon prolactin withdrawal were documented also in the 3D co-culture model of lactation using mammary epithelial and preadipocyte cell lines but, unluckily, control experiments with epithelial cells cultured without preadipocytes were not reported (Campbell et al., 2014). Thus, for the first time in organoid culture, we show that involution-like regression of epithelium occurs, at least in part, in an epithelium-intrinsic manner. Our observations do not contradict the crucial role of paracrine signaling between involuting epithelium and inflammatory stroma that is required for proper involution, including the leukemia inhibitory factor and transforming growth factor β signaling that activate STAT3 mediated regression of epithelium (Nguyen and Pollard, 2000; Kritikou et al., 2003; Hughes and Watson, 2012). Our results point to existence of epithelial-specific mechanisms of involution, for study of which our epithelial-only organoid model could be advantageous. However, to mimic involution fully, optimization of the culture conditions with cytokine cocktail would be required.

Several human diseases, developmental defects or insufficiencies in mammary epithelial tissue are linked to lactation and involution period. Among others, inadequate milk production affects many women after giving birth, especially in pre-mature deliveries (Olsen and Gordon, 1990; Kent et al., 2012). We propose that human breast tissue, gained from reduction mammoplasties, could be utilized to isolate primary human breast organoids and with some effort, findings from murine organoids could be translated into human organoids to study regulation of lactation using human tissue. Moreover, our organoid model could be used to investigate mechanisms of pregnancy-associated breast cancer, an aggressive form of breast cancer with peak of incidence within five years after delivery (Schedin, 2006). Mammary organoids isolated from genetic mouse models, such as animals carrying mutations in oncogenes or tumor suppressors, or organoids exposed to carcinogens could be employed in our lactation model to unveil mechanisms and signaling pathways leading to epithelial cell carcinogenesis.

## Author Contributions

JS, AC, EC and ZK performed the experimental work, AC, JS, ZK and HL contributed to experimental design and data analysis. AC, ZK and HL supervised the study. All the authors interpreted the data. ZK and HL acquired funding for the study. AC, JS and ZK wrote the manuscript. All authors discussed the results and approved the final version of the manuscript.

## Abbreviations

BOM: basal organoid medium;
Csn2: Casein2 - β-casein gene;
ECM: extracellular matrix;
FGF2: fibroblast growth factor 2;
LM: lactation medium;
Mmp: matrix metalloproteinase;
Wap: whey acidic protein.

## Acknowledgments

We are particularly grateful to Katarina Mareckova and Anas Rabata for their excellent technical support, and to Dr. Mari Aoki and Dr. Ulrich Boehm for providing cryosections of *Prlr-IRES-Cre;ROSA26-CAGS-GFP* mammary glands. We thank the Central Animal Facility of the Institut Pasteur and the Laboratory Animal Breeding and Experimental Facility of the Faculty of Medicine, Masaryk University. We also thank the Revive Consortium for funding the exchange program. We acknowledge the core facility CELLIM of CEITEC supported by the Czech-BioImaging large RI project (LM2015062 funded by MEYS CR) for their support with obtaining scientific data presented in this paper.

Work in the laboratory of HL is funded by Insitut Pasteur, Centre National pour la Recherche Scientificand the Agence Nationale de la Recherche (ANR-10-LABX-73; ANR-16-CE13-0017-01), Fondation ARC (PJA 20161205028, 20181208231) and AFM-Telethon Foundation. Work in the laboratory of ZK is funded by the Grant Agency of Masaryk University (grant no. MUNI/G/1446/2018). AC is funded by the postdoctoral fellowships from the Revive Consortium. JS was funded by the P-Pool (Masaryk University, Faculty of Medicine), Amgen Scholars Europe and Erasmus+ programs and by the Grant Agency of Masaryk University (grant no. MUNI/A/1565/2018). EC is funded by the PhD fellowship from Sorbonne Université.

## Contribution to the Field Statement

3D organoid systems are an indispensable tool for developmental morphogenesis studies. Primary mammary organoids, freshly isolated from murine tissue, keep most of the crucial architectural and functional in vivo features. Therefore, they have been widely employed to gain insights into mammary gland branching morphogenesis. However, later postnatal development of mammary gland, including pregnancy associated growth, lactation and involution, has been modeled in 3D scarcely and predominantly using established cell lines. Here, we describe a novel 3D culture model of mammary gland lactation and involution using primary mammary organoids, characterized in a comprehensive manner. In this model, primary mammary organoids first undergo branching morphogenesis, similarly to pregnancy in vivo. Then lactation, hallmarked by milk protein production and lipid synthesis, is induced and can last for at least 14 days. When the lactation stimuli are withdrawn, organoids enter involution-like regression. Importantly, our lactating organoids retain a contractile myoepithelial cell layer during long-term culture, marking the physiologically relevant epithelial architecture and functional state of lactating organoids. Therefore, our model can be employed by developmental biologists to study regulation of lactation and involution as well as by cancer researchers to study pregnancy-associated breast cancer.

