## Supplemental figures for "Primary mammary organoid model of lactation and involution"

Supplementary Material

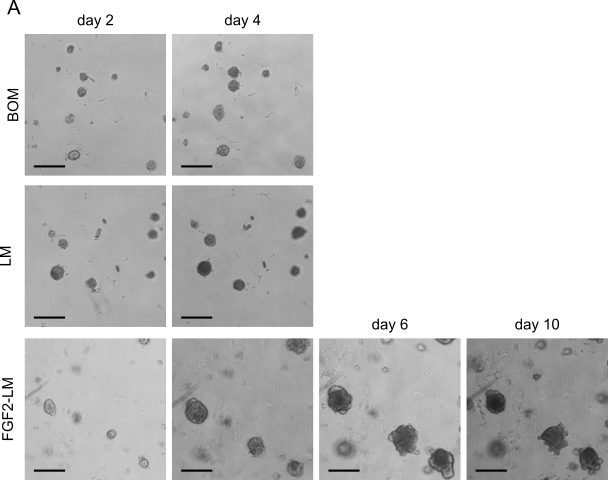

**Supplementary Figure 1.** **FGF2 pre-treatment enhances lactation – organoid morphology.** (A) Bright-field images of organoids treated with basal organoid medium (BOM), lactation medium (LM) or pre-treated with FGF2 and then treated with LM (FGF2-LM) at day 2, 4, 6 and 10. Scale bars represent 200 µm.

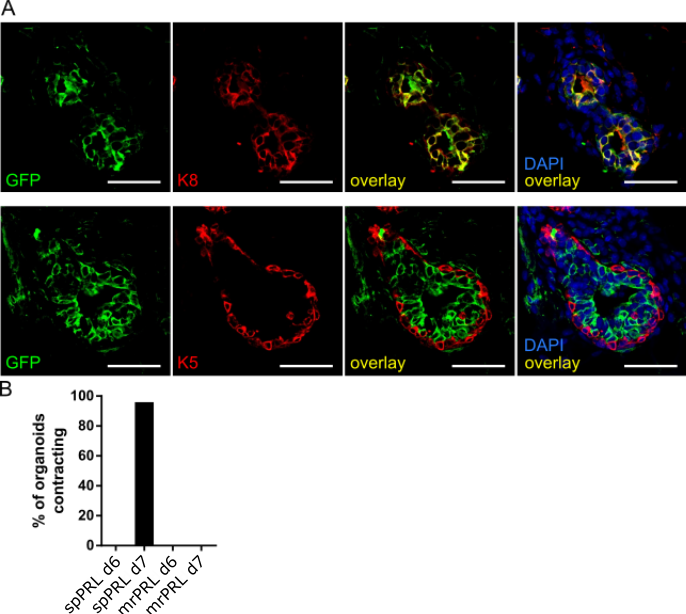

**Supplementary Figure 2.** **Contraction of organoids is not caused by direct prolactin signaling.** (A) Immunofluorescent staining of mammary gland from *Prlr-IRES-Cre;ROSA26-CAGS-GFP* mouse. Green, GFP in cells expressing prolactin receptor; red, keratin 5 (K5) or keratin 8 (K8); blue, DAPI). Scale bars represent 50 µm. (B) Quantification of contracting organoids from movies recorded on day 6 (before LM treatment) and day 7 (after LM treatment). Sheep pituitary (spPRL) or mouse recombinant prolactin (mrPRL) were used to prepare LM.
