## Supplementary material for "Primary mammary organoid model of lactation and involution": Movie 1, 2

### Slide 1
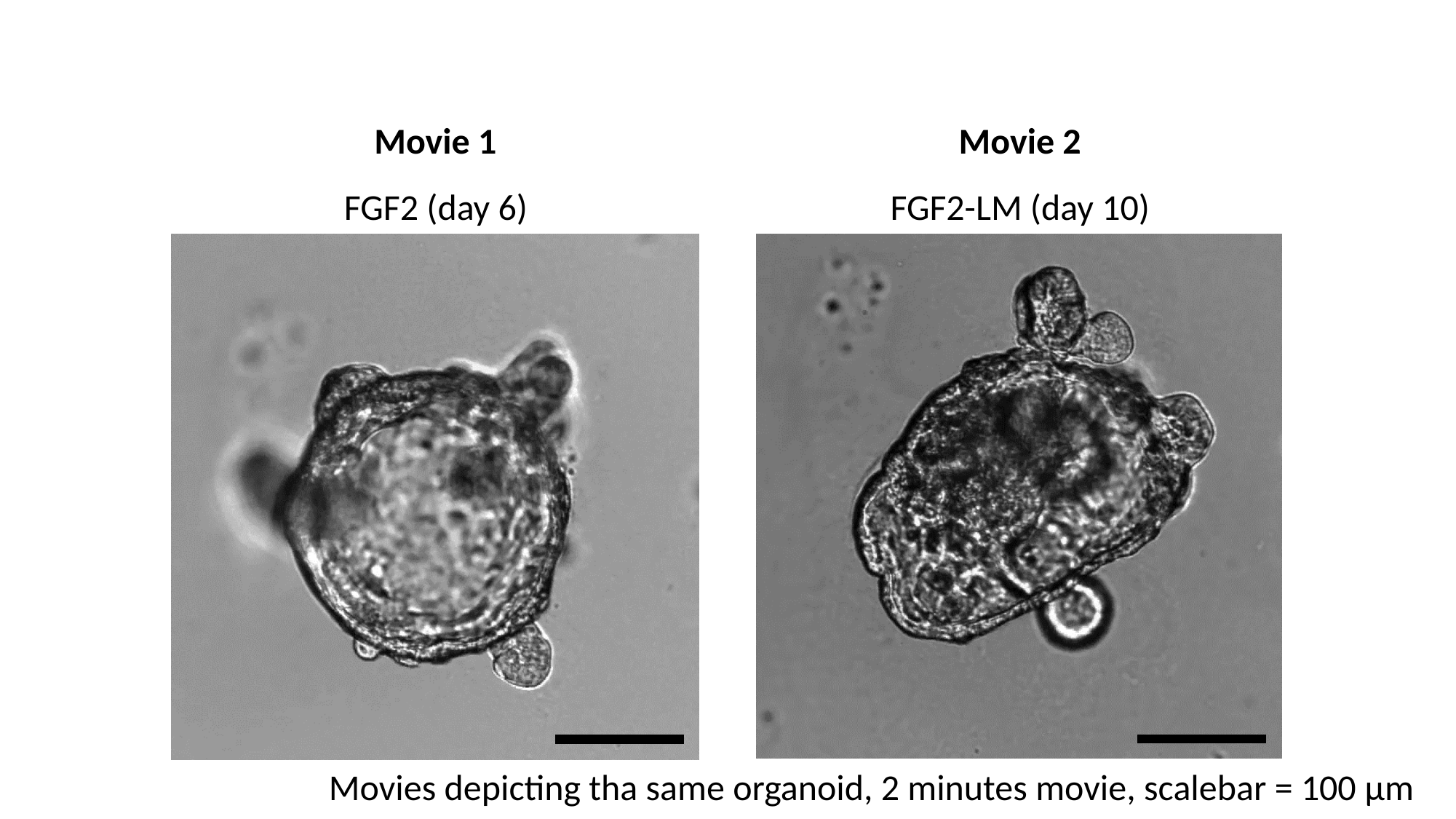

Movie 1
Movie 2
FGF2 (day 6)
FGF2-LM (day 10)
Movies depicting tha same organoid, 2 minutes movie, scalebar = 100 μm
